# Parallel loss of type VI secretion systems in two multi-drug resistant *Escherichia coli* lineages

**DOI:** 10.1101/2023.03.28.534550

**Authors:** Elizabeth A. Cummins, Robert A. Moran, Ann E. Snaith, Rebecca J. Hall, Chris H. Connor, Steven J. Dunn, Alan McNally

## Abstract

The repeated emergence of multi-drug resistant (MDR) *Escherichia coli* clones is a threat to public health globally. In recent work, drug resistant *E. coli* were shown to be capable of displacing commensal *E. coli* in the human gut. Given the rapid colonisation observed in travel studies, it is possible that the presence of a type VI secretion system (T6SS) may be responsible for the rapid competitive advantage of drug resistant *E. coli* clones. We employed large scale genomic approaches to investigate this hypothesis. First, we searched for T6SS genes across a curated dataset of over 20,000 genomes representing the full phylogenetic diversity of *E. coli*. This revealed large, non-phylogenetic variation in the presence of T6SS genes. No association was found between T6SS gene carriage and MDR lineages. However, multiple clades containing MDR clones have lost essential structural T6SS genes. We characterised the T6SS loci of ST410 and ST131 and identified specific recombination and insertion events responsible for the parallel loss of essential T6SS genes in two MDR clones.

**Data Summary:** The genome sequence data generated in this study is publicly available from NCBI under BioProject PRJNA943186, alongside a complete assembly in GenBank under accessions CP120633-CP120634. All other sequence data used in this paper has been taken from ENA with the appropriate accession numbers listed within the methods section. The *E. coli* genome data sets used in this work are from a previous publication, the details of which can be found in the corresponding supplementary data files 10.6084/m9.figshare.21360108 [1].

**Impact Statement:** *Escherichia coli* is a globally significant pathogen that causes the majority of urinary tract infections. Treatment of these infections is exacerbated by increasing levels of drug resistance. Pandemic multi-drug resistant (MDR) clones, such as ST131-C2/H30Rx, contribute significantly to global disease burden. MDR *E. coli* clones are able to colonise the human gut and displace the resident commensal *E. coli*. It is important to understand how this process occurs to better understand why these pathogens are so successful. Type VI secretion systems may be one of the antagonistic systems employed by *E. coli* in this process. Our findings provide the first detailed characterisation of the T6SS loci in ST410 and ST131 and shed light on events in the evolutionary pathways of the prominent MDR pathogens ST410-B4/H42RxC and ST131-C2/H30Rx.

## Introduction

*Escherichia coli* has been ranked globally as the number one causal pathogen of deaths associated with bacterial antimicrobial resistance (AMR) [2] and AMR has been declared by the World Health Organisation as a top ten global public health threat (www.who.int/news-room/fact-sheets/detail/antimicrobial-resistance). The emergence and evolution of multi-drug resistant (MDR) *E. coli* is a pressing and relevant issue for global healthcare as their extensive resistance profiles result in diminishing therapeutic options for treating infections.

A trait increasingly ascribed to MDR *E. coli* lineages is the ability to rapidly, and asymptomatically, colonise the human intestinal tract [3]. Multiple travel studies have now shown that people travelling from areas of low AMR incidence to AMR endemic regions become colonised by extended spectrum beta-lactam (ESBL)-resistant or MDR *E. coli* during travel [4; 5; 6]. Furthermore, genomic analysis has shown that the gain of MDR *E. coli* was due to the acquisition of a new MDR strain and not the preceding commensal *E. coli* becoming MDR [5]. The ability to displace and colonise may be attributed to the MDR phenotype itself, but longitudinal studies from the UK and Norway [7; 8] have shown that multi-drug resistance alone is not a sufficient driver for epidemiological success of *E. coli*. Recent metagenomic analysis has also revealed that colonisation by MDR *E. coli* does not disrupt the wider gut microbiome’s composition or diversity [9].

One possible reason for the ability of drug-resistant *E. coli* to displace resident commensal *E. coli* so rapidly is a result of the drug-resistant *E. coli* possessing a type VI secretion system that allows contact-dependent killing of the resident commensal *E. coli*. The type VI secretion system (T6SS) is a multi-functional apparatus that some Gram-negative bacteria possess to facilitate nutrient uptake [10], manipulation of host cells [11], and the killing of competing bacteria [12; 13; 14; 15]. T6SS distribution varies by environment [16; 17] and the presence of T6SSs varies on all taxonomic levels [16; 18; 19] with over-representation in Gammaproteobacteria [20]. T6SS genes can be gained and lost via horizontal gene transfer (HGT) [12; 18] and T6SS presence may be influenced by genomic incompatibilities, either between mobile genetic elements, donor and recipient bacteria, or via assembly of multiple T6SSs [16]. T6SSs are particularly prevalent in complex microbial communities that are host-associated [17; 18] and are commonly found in pathogens, including *E. coli* [16; 21; 22; 23].

*E. coli* is a relatively understudied organism within the research field of T6SSs, with many studies only focusing on specific pathotypes [22; 23; 24]. In *E. coli* T6SSs are classified into three distinct phylogenetic groups (T6SS-1 to T6SS-3) and all have been shown to be directly involved in pathogenesis and antibacterial activity [21]. The existing connections between T6SS and multi-drug resistance in other species [25; 26] makes T6SSs in *E. coli* an intriguing avenue of exploration in the context of pandemic MDR clones and how they become successful. Here, we assess the prevalence of T6SS genes across *E. coli* and identify no association with MDR or indeed any given lineages. We characterise the T6SS loci within MDR lineages ST410 and ST131 and show that successful MDR clones ST410-B4/H24RxC and ST131-C2/H30Rx have lost a functional T6SS through deletion and insertion events respectively.

## Methods

### Genome Collection

The 20,577 *E. coli* genomes used here were curated from EnteroBase [27] and were fully characterised and described in a previous study [1]. Briefly, they were chosen to represent the phylogenetic, genotypic and phenotypic diversity of the species. We include commensal, pathogenic, and generalist lineages and a spectrum of resistance profiles from susceptible to highly resistant. The data set is sampled from a variety of source niches. The assemblies covered six phylogroups and 21 different STs of *E. coli*; ST3, ST10, ST11, ST12, ST14, ST17, ST21, ST28, ST38, ST69, ST73, ST95, ST117, ST127, ST131, ST141, ST144, ST167, ST372, ST410, and ST648.

### T6SS database creation and interrogation

The SecReT6 [28; 29] experimentally validated database was retrieved and reformatted so that sequence identifiers were a suitable format for a custom ABRicate database. ABRicate (v0.9.8) was run using the default 75% minimum nucleotide sequence identity to obtain a list of T6SS genes that are present across our 21 *E. coli* sequence types. The results were summarised and partial hits (instances where a gene hit was split over multiple contigs) were accounted for and processed with a custom Python script (github.com/lillycummins/InterPangenome/blob/main/process_partial_hits.py). We chose to determine a gene as present in a genome if at least 85% of the gene was covered with a nucleotide sequence identity above the default 75% threshold. For the focused functional gene unit analysis, any matches to genes annotated as effector or immunity protein-encoding were removed from the search results.

### Genomic screening for insertion sequences and deletion events

Gene Construction Kit (v4.5.1) (Textco Biosoftware, Raleigh, USA) was used to annotate and visualise DNA sequences. Insertion sequences (ISs) were identified using the ISFinder database [30]. Target site duplications (TSDs) flanking ISs were identified manually. Two junction sequences specific for a given insertion were generated by taking 100 bp contiguous sequences that span the left and right ends of the insertion. Each junction sequence was comprised of 50 bp of the target IS and 50 bp of its adjacent sequence. To generate an ancestral sequence in silico, the interrupting IS and one copy of its associated TSD were removed manually. A 100 bp sequence that spanned the insertion point was taken to represent the naive site. A database of all ST131 genomes in the collection was generated for screening with standalone BLASTn [31]. The database was queried with insertion-junction sequences and the naive site sequence to determine whether an insertion was present, with only complete and identical matches to the 100 bp query sequences considered positive matches. The same 100 bp indicator sequence method was used to identify deletion events.

### Genome sequencing

To the best of our knowledge, no complete reference genome for ST131-A/H41 was publicly available so we generated sequence data for the strain USVAST219 [32]. DNA was extracted using the Monarch Genomic DNA Extraction Kit (New England Biolabs, Massachusetts, USA) before sequencing with MinION (Oxford Nanopore Technology, UK) using R9.4.1 flow cells. The data was basecalled with Guppy (v6.0.1) (github.com/nanoporetech/pyguppyclient) and adapters were trimmed with qcat (v1.1.0) (github.com/nanoporetech/qcat). Short-read genome sequencing was provided by MicrobesNG (microbesng.com) where DNA was prepared using the Nextera XT kit (Illumina, San Diego, CA, USA) and sequenced on the Illumina HiSeq platform (Illumina). Long and short-read data are both available within BioProject PRJNA943186. A hybrid assembly using both the long and short-read sequencing data was created with Unicycler (v0.4.8) [33].

## Results

### T6SS gene presence varies between and within phylogroups and sequence types

All 20,577 *E. coli* genomes interrogated contained at least one T6SS gene from the SecReT6 experimental database [28; 29]. There was variation in the average number of T6SS genes present per ST and the average number of genes per ST within phylogroups (Figure 1). ST131 exhibited the largest range (2 to 29) of T6SS genes, highest number of T6SS genes in a single genome (n = 29), and highest average number of T6SS genes (joint with ST12, ST14, ST17, and ST127). The distribution of the number of T6SS genes present per genome did not align with phylogroups. STs known to contain MDR lineages, ST131, ST167, ST410, and ST648, did not possess a similar range or average of T6SS genes (Figure 1). Grouping genes into functional units is therefore necessary to gain insight into how many potentially functional T6SSs are present within STs and whether this may correlate with multi-drug resistance.

**Fig. 1.**
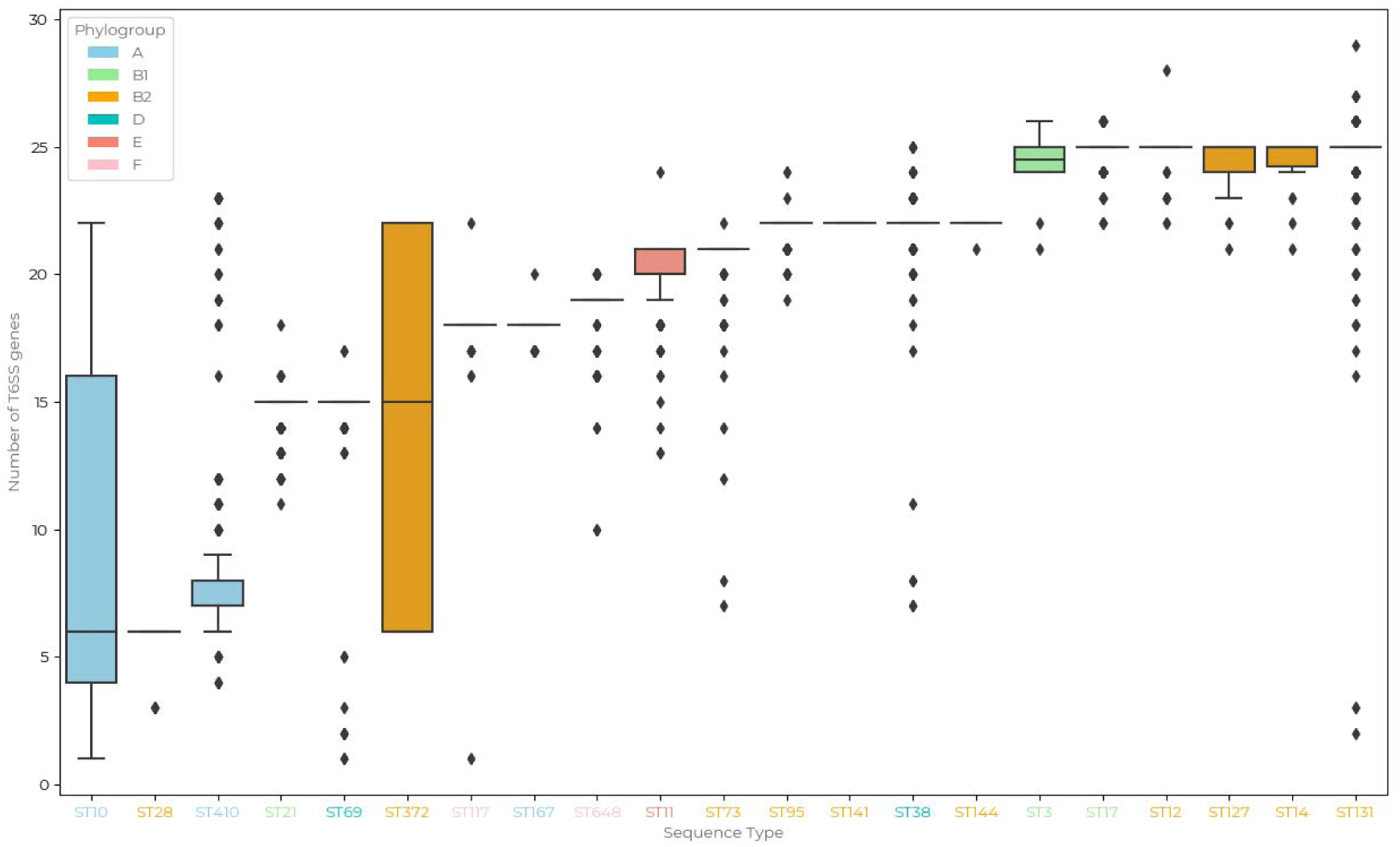
Distributions of the number *of type VI secretion system (T6SS) genes from the SecReT6 experimentally validated database [28; 29] across 21 sequence types of E. coli shows gene carriage varies by sequence type and phylogroup. Sequence types are colour coded according to their respective phylogroups: orange = A; green = B1; purple = B2; light blue = D; blue = E; red = F. A gene is determined as* present if it exceeds a DNA sequence identity of 75% and a coverage of 85%.

### Presence of structural T6SS genes in lineages containing MDR clones

We examined T6SS gene units in STs containing MDR clones in greater detail. The MDR clones ST131-C2/H30Rx and ST410-B4/H24RxC were selected for further investigation due to their global distribution and clinical burden [7; 8; 34; 35]. Figure 2 displays the presence of structural T6SS genes within ST410 (Figure 2a) and ST131 (Figure 2b) with the corresponding clades highlighted on the phylogenies. Structural genes were the focus of this search because they are well characterised and vary less in comparison to effector/immunity proteins [21]. In both STs, we found a correlation between clades possessing MDR clones and the absence of structural T6SS genes (Figure 2).

**Fig. 2.**
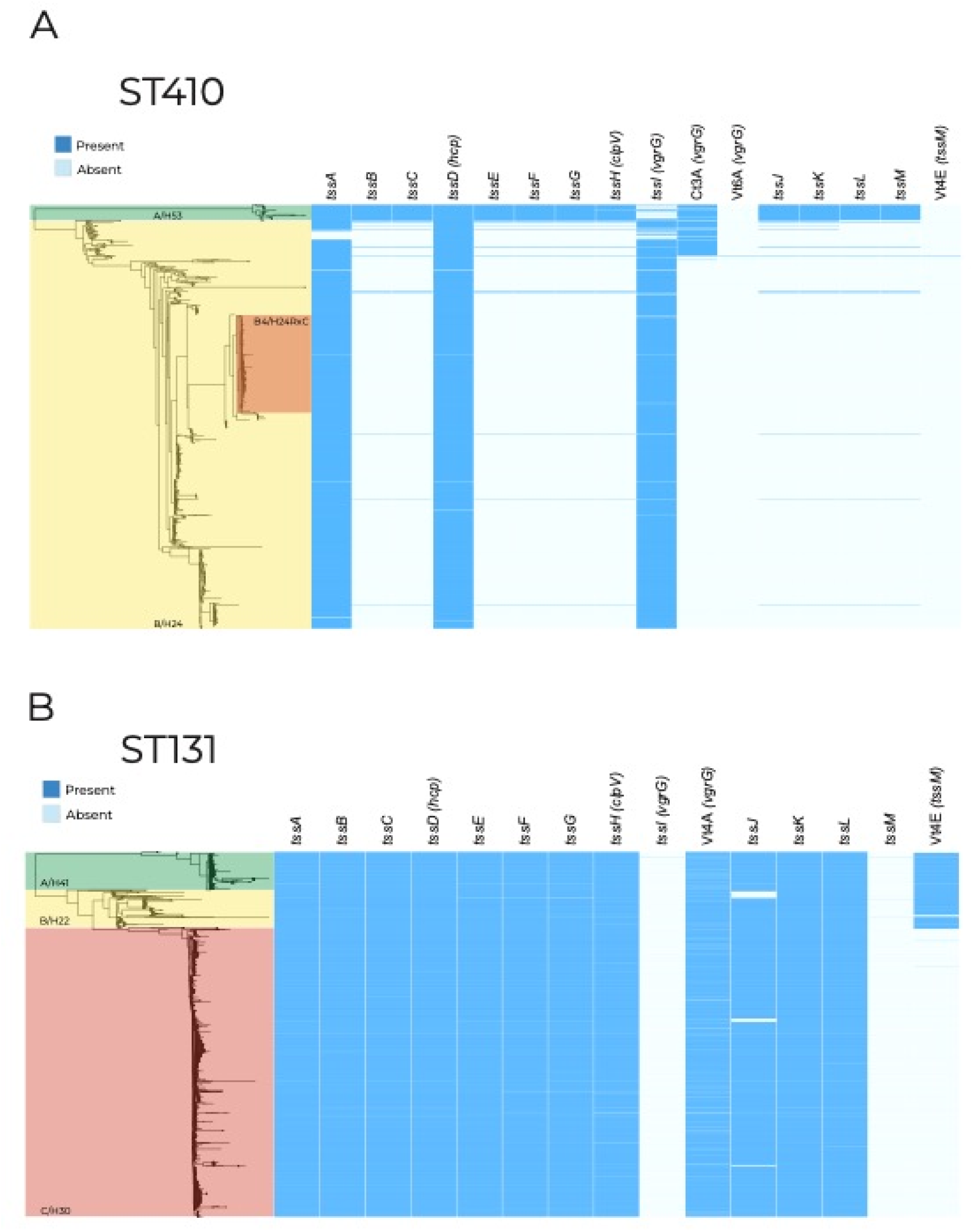
Presence of structural experimentally validated type VI secretion genes taken from the SecReT6 database [28; 29] across (A) 1,006 ST410 genomes and (B) 3,186 ST131 genomes. Phylogenetic clades are labelled clade name/fimH allele.

### Presence of structural T6SS genes in ST410

Within ST410, *tssABCDEFGHIJKLM*, (henceforth referred to as the *tss* region) was present in 94.59% (n = 34) of clade A/H53 (Figure 2a). The organisation of genes in this region classifies the system as T6SS-2. Of the three genomes that did not contain a full *tss* region, one genome was only missing *tssH* due to poor sequence coverage (coverage <10x for the three contigs spanning *tssH*). In the two other genomes only *vgrG* was missing, caused by a combination of low coverage and the presence of multiple *vgrG* genes, which is known to lead to sequence fragmentation. We conclude that the native T6SS of ST410-A/H53 is conserved across the clade.

The *tss* region was present in just 0.83% (n = 8) of clade B/H24 genomes. Two ST410-B/H24 genomes did not contain *tssI/vgrG* and four genomes did not contain *tssLM*, but these otherwise contained all *tss* region genes. While isolates from clade A/H53 are largely drug susceptible, clade B/H24 contains the globally distributed multi-drug resistant clone ST410-B4/H24RxC, which is highlighted in Figure 2a [34; 36]. In the eight rare instances where a full *tss* region was present within the B/H24 clade, five genomes contained *tss* regions most closely related (≥98% nucleotide identity) to that of the T6SS-2 region of F-type plasmid pSTEC299_1 (GenBank accession: CP022280) when comparing to the NCBI non-redundant database. Two of the remaining three genomes contained *tss* regions that were closely related (≥98% nucleotide identity) to chromosomal T6SS-1 regions from other *E. coli* (GenBank accessions: CP062901, CP091020). These findings together suggest that, although rare, multiple acquisitions of distinct *tss* regions have occurred within ST410 clade B/H24.

### Characterisation of the ST410 type VI secretion locus

We resolved the structure of the complete 28.3 kb T6SS locus of ST410-A/H53 using a draft ST410-A/H53 assembly (EnteroBase assembly barcode: LB4500AA_AS), partially scaffolded (Fig. S1) using the complete genome of ST88 *E. coli* RHBSTW-00313 (GenBank accession: CP056618). ST88 belongs to the same clonal complex as ST410 (CC23) and was used because there is no complete reference genome for ST410-A/H53 and we were unable to access a strain to generate a closed genome. The complete T6SS region contained 18 open reading frames (ORFs), 16 of which could be assigned functions (Figure 3). Among these were representatives of all 13 *tss* genes required for synthesis of a type VI secretion apparatus (*tssA* to *tssM*), along with two additional different *tssA* and *tssD* genes. The recombination hotspot *rhsG* gene, which is known to encode a toxin [39], is located downstream of *vgrG* and is the likely determinant for this systems’ effector protein. The locus also contains a gene for an FHA-domain containing protein, which is known to be involved in T6SS activation in *Pseudomonas aeruginosa* [40]. The structure of this region is indicative of a T6SS-2 type system [21] and we conclude that this region of the ST410-A/H53 chromosome includes all determinants necessary for the production of a functional T6SS.

**Fig. 3.**
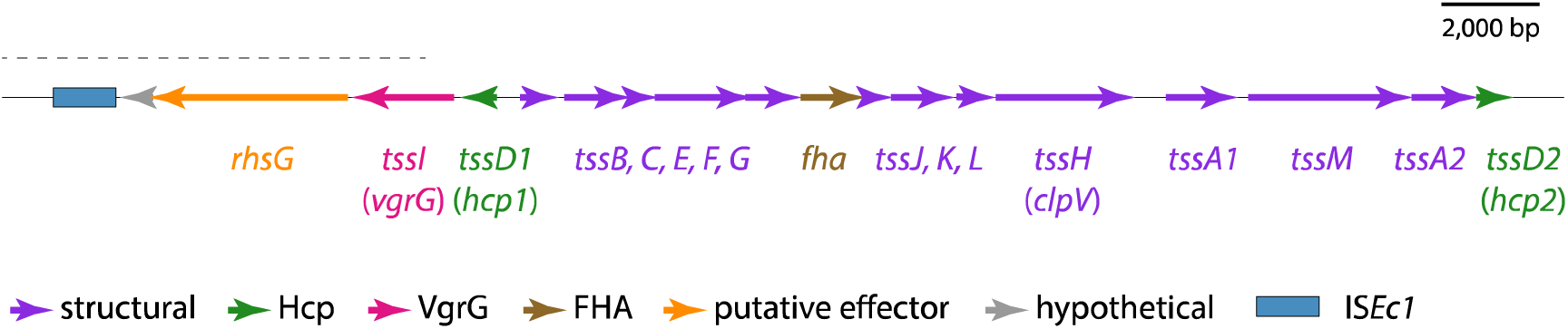
The ST410-A/H53 chromosomal T6SS locus. The extents and orientations of open reading frames are indicated by labelled arrows, with colours corresponding to protein types as outlined in the key below. The part of the sequence for which sequence from ST88 was used as a scaffold is indicated by the dotted black line above. FHA: forkhead-associated. Drawn to scale from EnteroBase assembly barcode: LB4500AA_AS and GenBank accession: CP056618.

### Deletion events account for the loss of T6SS in ST410-B/H24

To explain the absence of T6SS genes in ST410-B/H24, we annotated the chromosomal region of reference strain SCEC020026 (GenBank accession CP056618) that corresponds to where the *tss* region characterised in ST410-A/H53 is located. Adjacent to the *tssA2* and *tssD2* genes, which are present in the majority of ST410-B/H24 genomes (Figure 4a), we found remnants of *rhsG* and *tssM* (Figure 4b). Relative to the complete *tss* region in clade A/H53, clade B/H24 has lost 21.0 kb between *rhsG* and *tssM* in a deletion event. As there are no mobile genetic elements present at the deletion site, which is located within a recombination hotspot, the deletion event was most likely the result of recombination. To determine whether this deletion event was responsible for the loss of the *tss* region in other ST410-B/H24 isolates, we queried their genomes with the unique 100 bp sequence found at the *rhsG-tssM* junction in SCEC020026. The junction sequence (G-M) was present in the majority of ST410-B/H24 genomes (n = 941 out of 969; Figure 4a), indicating that this deletion event occurred as a single event early in the evolution of ST410-B/H24 and not as multiple independent events within this clade.

**Fig 4.**
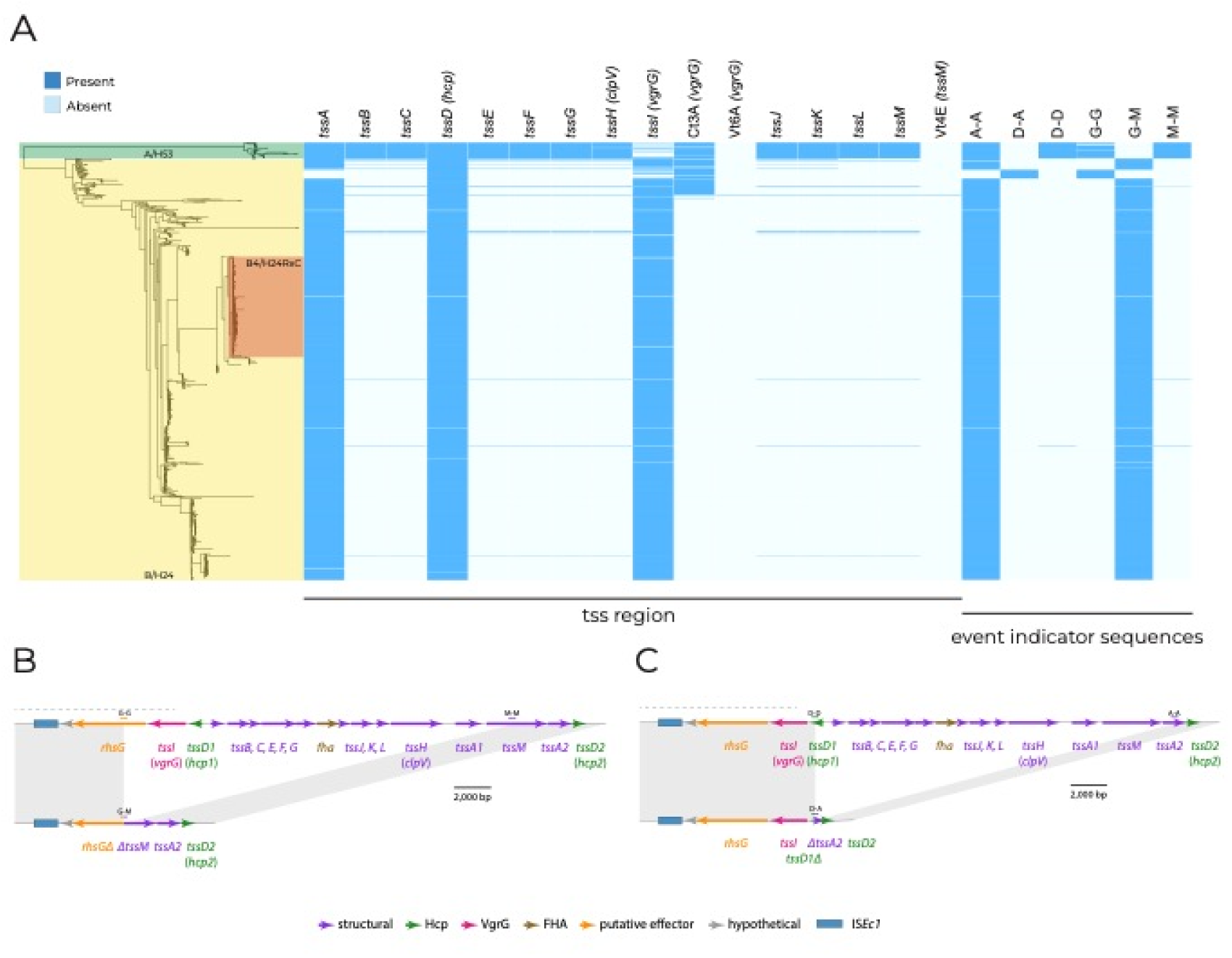
(A) ST410 phylogeny. Genomes are shaded according to their clade and fimH allele. The presence of tss region genes and 100bp deletion event indicator sequences is shown by blue shading to the right of the phylogeny. (B) Deletion of 21.0 kb segment of tss region between rhsG and tssM. (C) Deletion of 19.8 kb segment of tss region between tssD1 and tssA2. Chromosomal sequence is shown as thin black lines, with open reading frames indicated by arrows beneath. Dotted black lines indicate sections of region where sequence from ST88 was used as a scaffold.

Of the 28 ST410-B/H24 genomes that did not contain the *rhsG-tssM* junction sequence, 20 were clustered in the phylogeny and also lacked *tssA* (Figure 4a). This suggests that a different or additional deletion event may have occurred in a small sub-clade of ST410-B/H24. To determine this, we annotated a contig from one of these 20 genomes (EnteroBase assembly barcode: ESC_HA8479AA_AS), which contained *tssD2* and exhibited a different *tss* region configuration. To confirm the structure of this region, we used the ESC_HA8479AA_AS *tss* contig to query GenBank and found an identical sequence in a complete ST410 genome (strain E94, accession: CP199740) that had been published after our data set was assembled. We then used the E94 genome as a representative for the 20 clustered genomes. This revealed the absence of a 19.8 kb segment of the *tss* region between *tssD1* and *tssA2* (Figure 4c). The presence of the same *tssD-tssA* junction sequence (D-A) in all 20 clustered genomes indicated that this deletion event was responsible for the absence of *tss* genes in this sub-clade of ST410-B/H24.

### Presence of structural T6SS genes in ST131

Within ST131, the ancestral and less drug-resistant clades A/H41 and B/H22 possess a gene labelled Vt4E from the SecReT6 database, which is not present in clade C/H30 (Figure 2b). To deduce the function of Vt4E, we used BLASTn to search [31] the GenBank non-redundant nucleotide database, which revealed that Vt4E shared 100% sequence identity with *vasK*. The *vasK* gene is a homologue of *tssM/icmF* [37; 38] which is a component of the inner membrane complex of T6SSs [21]. The absence of a functional VasK protein has been shown to render the T6SS non-functional in *Vibrio cholerae* [37].

### Characterisation of the ST131 type VI secretion locus

To facilitate comparative analyses, we characterised the T6SS-determining region of the ancestral ST131 lineage, ST131-A/H41. First, we mapped the complete and uninterrupted T6SS locus found in a 39,211 bp region of the USVAST219 chromosome (GenBank accession: CP120633) which was flanked by copies of a perfect 41 bp repeat sequence (Figure 5). The region contained 27 ORFs, 22 of which could be assigned functions (Figure 5). Among these were representatives of all 13 *tss* genes required for synthesis of a type VI secretion apparatus (*tssA* to *tssM*), along with the structural gene tagL and a second, different *tssA* gene. Genes for Hcp (*tssD*) and PAAR-domain containing proteins, along with three different *vgrG* (*tssI*) genes, encode the core and spike of the secretion apparatus in ST131-A/H41. Downstream of each *vgrG*, we found ORFs that encode putative T6SS effector proteins; a M23-family peptidase, a lytic transglycosylase and proteins with DUF4123 and DUF2235 domains, which have been associated with T6SS effectors [41]. This aligns with prior research by Ma and colleagues that identified diverse effector/immunity modules within *vgrG* modules [38]. Together, the presence and order of these components classify this locus as encoding a T6SS-1 type system [21]. We conclude that this region of the ST131 chromosome likely includes all determinants sufficient for production of a functional T6SS.

**Fig. 5.**
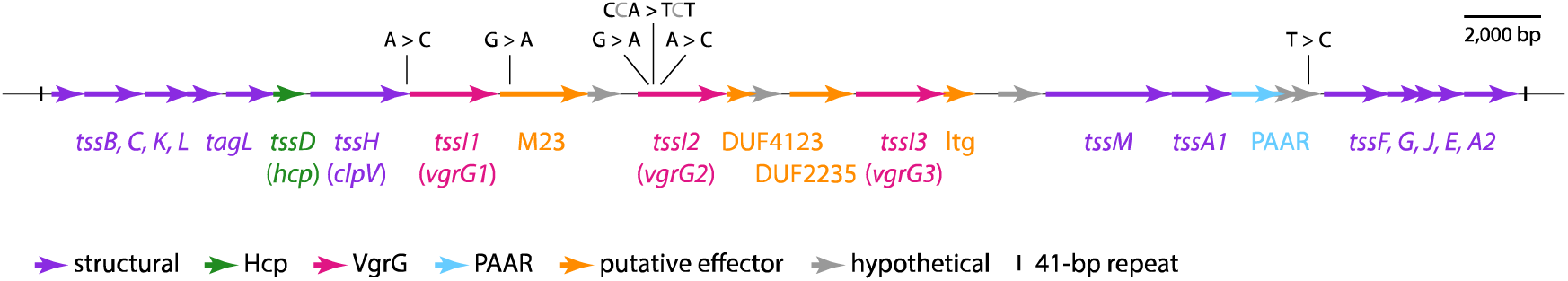
The ST131 chromosomal T6SS locus. The extents and orientations of open reading frames are indicated by labelled arrows, with colours corresponding to protein types as outlined in the key below. SNPs that differentiate clade C strain EC598 from clade A strain USVAST219 are shown in black text above. M23: M23-family peptidase, ltg: lytictransglycosylase. Drawn to scale from GenBank accession CP120633.

Reference strain EC958 (GenBank accession: HG941718) [42] was used to represent ST131-C2/H30Rx in comparisons with ST131-A/H41. The T6SS regions of USVAST219 and EC958 were almost identical, differing by just seven single nucleotide polymorphisms (SNPs) (shown in Figure 5). The only SNP located in a structural gene (*tssH*) was silent. We therefore conclude that this chromosomal region encodes the native T6SS of ST131, and is conserved in clades A/H41, B/H22, and C/H30, consistent with the presence/absence data shown in Figure 2b. However, the EC958 T6SS region does not encode TssM.

### tssM is interrupted by ISEc12 in ST131-C/H30

To determine the cause of *tssM* loss in ST131-C/H30, we examined the complete genome of reference strain EC958 [42] in further detail. We found that *tssM* had been interrupted by a copy of IS*Ec12*, which was flanked by the 5 bp target site duplication ACTGC (Figure 6). The *tssM* reading frame was split approximately in half by the insertion, with 1,570 bp located to the left of IS*Ec12*, and 1,768 bp to the right. To deduce whether other *tssM* homologues were present in EC958, and could encode replacements for the interrupted *tssM*, we used tBLASTn to query the complete EC958 chromosome with the TssM sequence encoded by USVAST219.

**Fig. 6.**
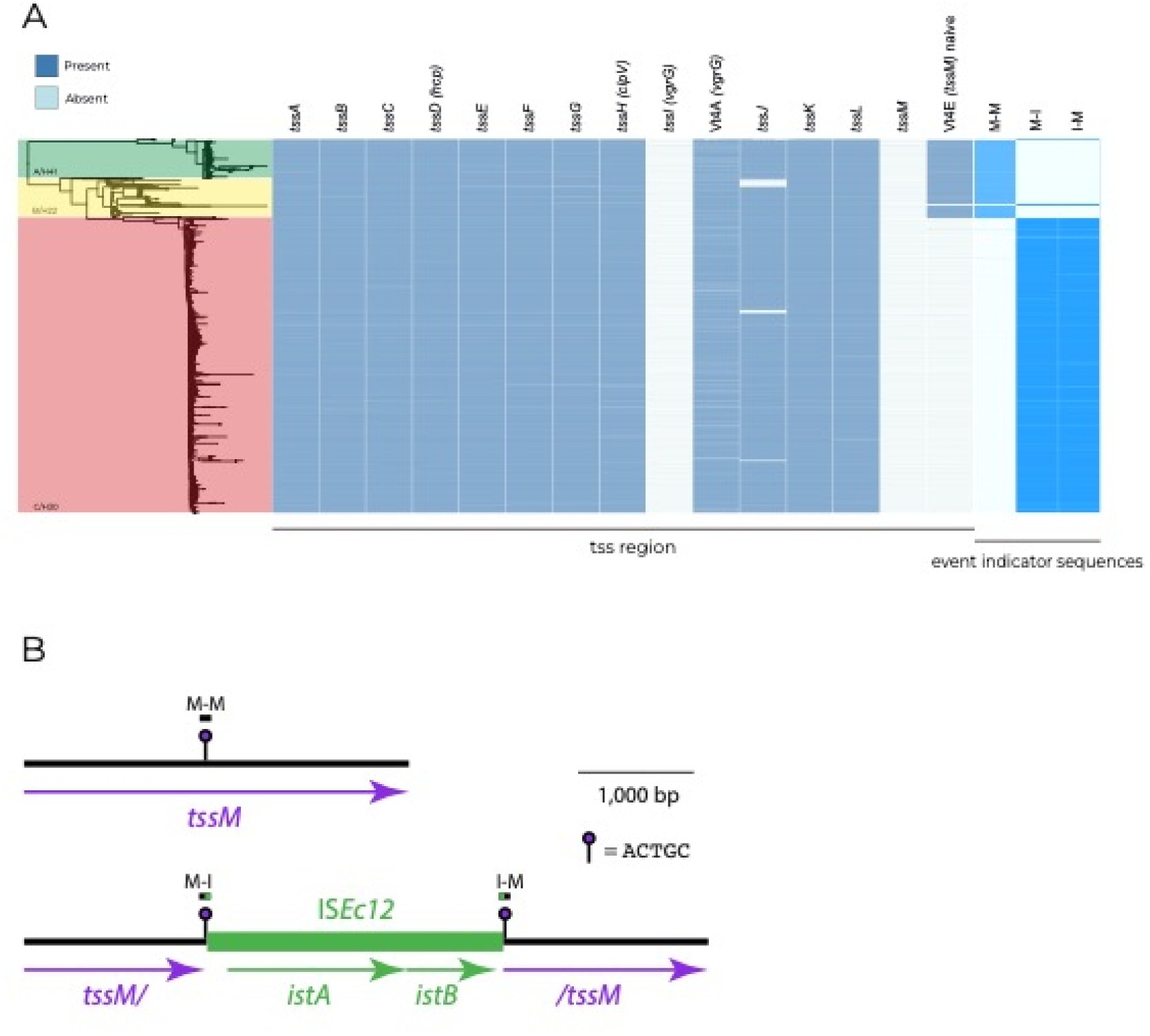
(A) ST131 phylogeny. Genomes are shaded according to their clade and fimH allele. The presence of the tss region and 100 bp ISEc12 insertion event indicator sequences is dictated by blue shading to the right of the phylogeny. (B) Interruption of tssM by ISEc12. Chromosomal sequence is shown as a thin black line, and ISEc12 as a thicker green line, with open reading frames indicated by arrows beneath. The positions of target site duplication sequences and extents of indicator sequences are shown above. Drawn to scale using sequence obtained from GenBank accessions: CP120633 and HG94718.

This query did not return any hits, apart from the fragmented sequences produced by the IS*Ec12*-interrupted *tssM*. It has been proven that the deletion of *tssM* prevents T6SS activity in *E. coli* [38] and the importance of *tssM* homologues for T6SS functionality in other species is well described [43; 44]. The IS*Ec12* insertion has almost certainly rendered *tssM* non-functional in EC958, and in the absence of *tssM* redundancy, it appears most likely that the entire T6SS itself is also non-functional.

From the EC958 genome, we generated 100 bp sequences that span the left and right ends of the IS*Ec12* insertion (marked M-I and I-M in Figure 6b), and from the USVAST219 genome a 100 bp sequence that represents an ancestral *tssM* uninterrupted by this insertion (marked M-M in Figure 6b). To determine whether the same insertion event was responsible for the absence of *tssM* in all clade C/H30 genomes, we screened the ST131 collection for the three 100 bp sequences. The naive M-M sequence was absent in just 2.97% (n = 21) of clade A/H41 and B/H22 genomes, confirming that almost all of them contained uninterrupted *tssM* (Figure 6a). 2,462 out of the 2,478 (99.48%) clade C/H30 genomes lacked the M-M sequence but contained one or both of the left and right IS*Ec12* junction sequences M-I and I-M (Figure 3). Of the 16 remaining clade C/H30 genomes, three genomes contained M-M, and 13 genomes did not include either M-M or either of the IS*Ec12* junction sequences (Figure 6a). Manual inspection of the 13 genomes that lacked indicator sequences revealed that in six the IS*Ec12* insertion was present, but low assembly quality had resulted in truncated contigs that were too short for recognition with the strict threshold used for junction sequence detection. The remaining seven genomes were excluded from further analysis, as low coverage appears to have resulted in highly-fragmented assemblies such that the structure of the T6SS region was impossible to determine accurately. Thus, while *tssM* is uninterrupted in 97.03% (n = 687) of ST131 clade A/H41 and B/H22 genomes, it has been interrupted by IS*Ec12* at exactly the same position in 99.78% (n = 2465) of clade C/H30 genomes. We conclude that the loss of *tssM* in ST131-C/H30 was the result of a single IS*Ec12* insertion event that occurred in an ancestral strain.

### The tss region is conserved in other phylogroup B2 lineages

Other pathogenic lineages of *E. coli* were examined to investigate whether an incomplete *tss* region was specific to clades containing MDR clones or a general feature of successful extraintestinal pathogenic *E. coli* (ExPEC). ST73 and ST95 were selected as comparators due to their documented prevalence in clinical settings in the UK [7; 45]. They and ST131 belong to phylogroup B2, which would also allow us to address phylogenetic signal influencing the presence of the *tss* region. The *tss* region was found to be consistently present in both lineages (Figure 7a and b), demonstrating that T6SS presence varies between pathogenic *E. coli* lineages within phylogroup B2.

**Fig. 7.**
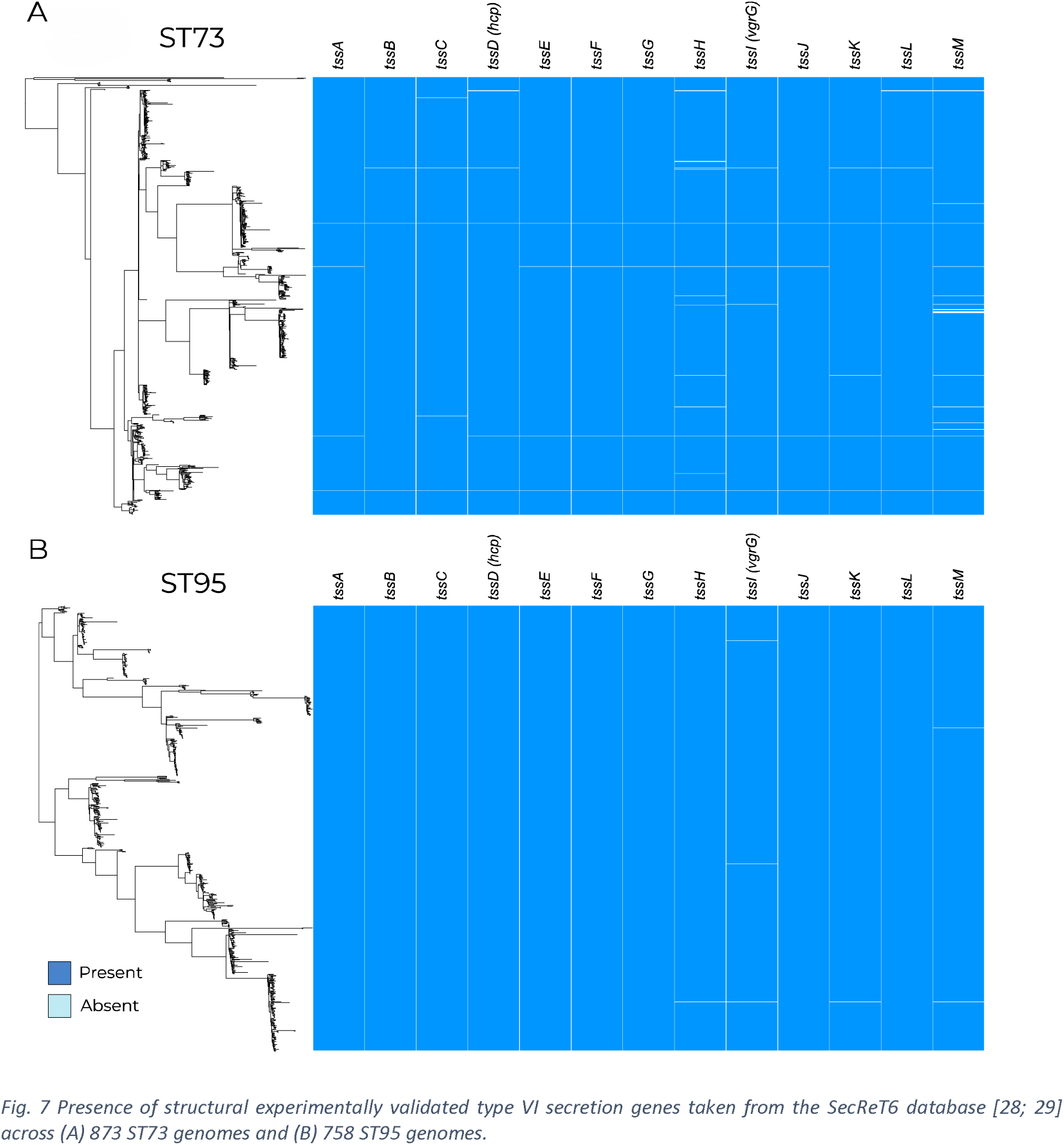
Presence of structural experimentally validated type VI secretion genes taken from the SecReT6 database [28; 29] across (A) 873 ST73 genomes and (B) 758 ST95 genomes.

## Discussion

Extensive work has been done to characterise the T6SS in terms of its structure, function, variability and prevalence in various genera. However, this is the first study, to the best of our knowledge, to determine the prevalence of the *tss* region within MDR ExPEC lineages. We have particularly focused on ST410 and ST131 as they are known to cause infections worldwide [34; 35; 36; 46]. We have identified the evolutionary events that led to the loss of key structural T6SS components in these lineages via recombination or insertion events. Our study has yielded new insights on the evolution of T6SS functionality in two of the most clinically-relevant MDR *E. coli* lineages, and has provided a broader overview of T6SS determinants in other lineages.

In both ST410 and ST131, we have observed a loss of T6SS that appears to coincide with the acquisition of MDR. This suggests we may have uncovered a common evolutionary trajectory in two MDR lineages. We have previously described the important role of potentiating mutations involved in colonisation factor determinants, anaerobic metabolism genes, and intergenic regions as being common evolutionary trajectories in MDR clone formation [47]. The loss or degradation of a T6SS could perhaps integrate into the stepwise evolutionary course of ST410-B4/H24RxC and ST131-C2/H30Rx. The structural components for a complete T6SS were chromosomal and conserved within the ancestral clades ST410-A/H53, ST131-A/H41 and ST131-B/H22. We found that the T6SS in ST410-B/H24 was inactivated by deletion events and that the T6SS in ST131-C/H30 was inactivated by an insertion event. In ST131, the rare instances where *tssM* is interrupted in clade A/H41 and B/H22, and uninterrupted in clade C/H30 (Figure 6a), might be explained by inter-lineage horizontal gene transfer and homologous recombination, but the T6SS region is too well-conserved (>99.99% ID between lineages) to accurately detect recombinant sequences. In both ST131 and ST410, the inactivation of T6SS occurring prior to or during the formation of MDR clades implies that these events may have contributed to their successful emergence and expansion, but we cannot generalise to all MDR lineages as further analyses of the common MDR lineages ST167 and ST648, not shown here, showed intact T6SS regions.

It has previously been suggested that acquiring MDR plasmids is a vital step in the evolution of pandemic clones [47]. We speculate that the lack of a functional T6SS allowed both ST410-B/H24 and ST131-C/H30 to become more receptive to cell-to-cell contact, and therefore conjugative transfer. The deleterious effect of T6SS on plasmid conjugation has been demonstrated in *Acinetobacter baumannii*, where a conjugative plasmid has been shown to repress the host T6SS in order to increase conjugation rates [26]. While the *A. baumannii* experiments involved inhibition of T6SS in donor cells, the absence of T6SS in potential recipient cells seems likely to increase their permissiveness to plasmid transfer. The acquisition of conjugative MDR plasmids by ST410-B/H24 and ST131-C/H30 might therefore have been facilitated by their loss of functional T6SS.

Our genomic analyses, combined with existing literature, suggest that the MDR clone ST131-C2/H30Rx does not have a functional T6SS due to the interruption of a single gene, *tssM*. However, the functionality of the uninterrupted *tss* region in the ancestral ST131 clades A/H41 and B/H22 has not been verified experimentally. Further work to determine the functionality of the T6SS in all three ST131 clades is therefore required. Final experimental validation would require complementation of *tssM* to restore function of the T6SS in ST131-C/H30.

The data presented here does not support the hypothesis that the ability of drug resistant *E. coli* to displace resident commensal *E. coli* is due to the production of a T6SS. Phage, colicins, or other diffusible elements are alternative explanations that should be considered. By focusing on T6SS, we have uncovered parallel evolutionary outcomes in ST410-B4/H24RxC and ST131-C2/H30Rx, where T6SS determinants have been lost by MDR clones. Our work suggests that the loss of functional T6SS might have played a role in the evolution and success of the pandemic MDR *E. coli* clones in ST131 and ST410.

## Author Statements

### Author Contributions

E.A.C.: data curation, methodology, formal analysis, visualisation, writing – original draft preparation and review and editing. R.A.M.: methodology, formal analysis, writing – original draft preparation and review and editing. A.E.S: methodology. R.J.H.: writing – review and editing. C.H.C.: data curation. S.J.D.: data curation. A.M.: conceptualisation, supervision, writing – review and editing.

## Acknowledgements

We would like to thank Professor Jose Bengoechea for helpful discussion and all members of the McNally group for their support.

