## Supplementary figures for "Parallel loss of type VI secretion systems in two multi-drug resistant *Escherichia coli* lineages"

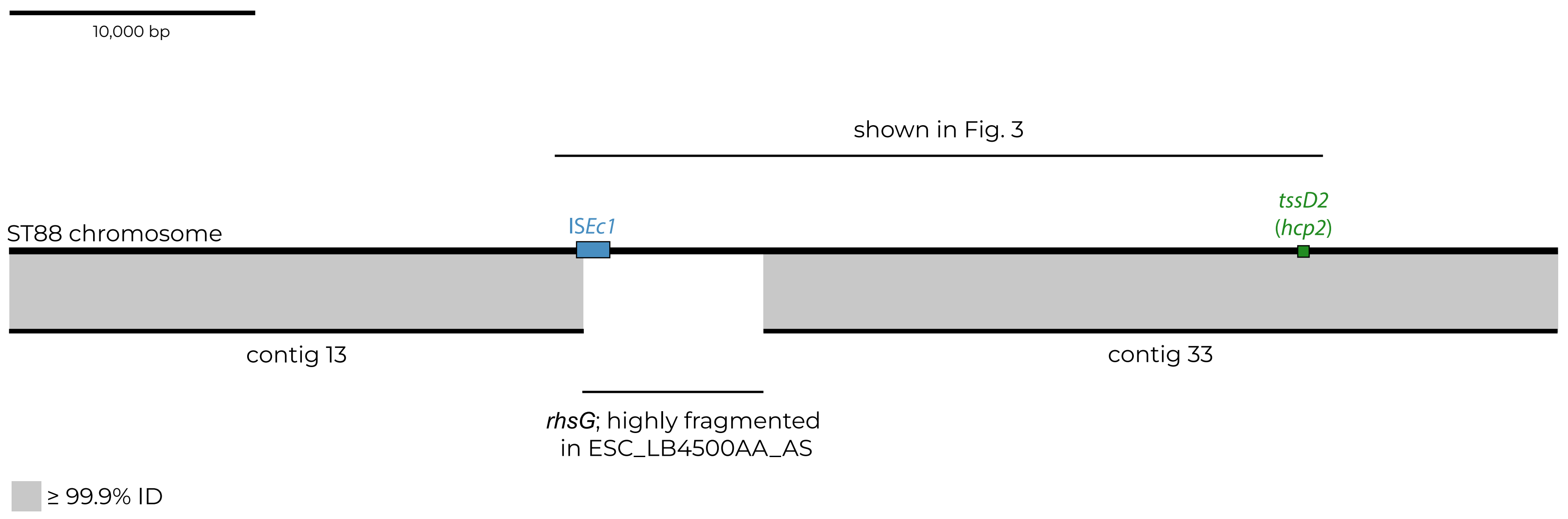


**S 1.** Scaled schematic showing the alignment of tss region sequences in the draft ST410 genome (EnteroBase assembly barcode LB4500AA_AS and the complete ST88 genome (GenBank accession CP056618) that was used as a scaffold when determining the structure of the region in ST410 clade A. Horizontal lines represent sequences, connected by grey shading indicative of alignments with $\geq$ 99.9% nucleotide identity. Contig numbers in LB4500AA_AS are indicated below, and the highly fragmented 'recombination hot spot' region is labelled. The boundaries of the tss region are marked with IS*Ec1* on the left and *tssD2* on the right, and the extent of the sequence shown in Figure 3 is indicated above.
